# Novel Cardiometabolic Factors Regulate Neurite Outgrowth in Cancer Chemotherapy-Induced Cardiotoxicity

**DOI:** 10.64898/2026.06.12.731794

**Authors:** Olivia Sloan, Skylar Orr, Viswanathan Rajagopalan

**Author notes:** Address for Correspondence: Viswanathan Rajagopalan, Ph.D., FAHA, FCVS, Associate Professor, New York Institute of Technology College of Osteopathic Medicine at A-State, 119 State University, Jonesboro, AR 72467.

## Abstract

**Background:** Cardiovascular diseases and cancer are the leading causes of death in the United States and worldwide. Although various therapies against cancer improve patient survival, cardiotoxicity remains a life-threatening adverse outcome, with emerging evidence of downstream effects, including neural dysfunction. While autonomic regulation of the cardiovascular system is well-studied, regulation of the nervous system by the heart is not fully clear. We hypothesized that cardiac cells secrete non-canonical paracrine metabolic factors that support neuronal growth and function, and chemotherapy disrupts this signaling.

**Methods:** We employed co-culture models of the well-established H9C2 cardiac and PC12 neuronal cell lines and human induced pluripotent stem cells (hiPSCs), and assessed them with molecular, omic, biochemical, morphological, physiological, and pharmacological assays.

**Results:** Healthy H9C2 cells robustly induced PC12 neurite outgrowth (neurite length and number of neurite-bearing cells) both directly (with cellular contact) and indirectly (only conditioned media), whereas doxorubicin-exposed H9C2 cells failed to produce this effect. Recently approved anti-cancer agents (2020 or later) also reduced or attenuated cardiac cell-induced outgrowth. Untargeted metabolomic analysis of conditioned media revealed multiple novel potential neurite-promoting factors, and pharmacologically inhibiting them significantly reduced PC12 neurite outgrowth. The analysis also identified distinct metabolites that were differentially regulated following doxorubicin exposure. These findings were further supported in a hiPSC-based model, in which conditioned media from doxorubicin-injured hiPSC cardiomyocytes reduced βIII-tubulin intensity and norepinephrine secretion in hiPSC-derived sympathetic neurons.

**Conclusion:** Together, these findings unravel a new line of research on cardio-neuronal communication and reveal novel metabolic targets that may inform future strategies to mitigate neurotoxicity induced by chemotherapy-associated cardiac injury.

## INTRODUCTION

Cardiovascular diseases and cancer are the leading causes of mortality in the United States and across the world [1–3]. Patients with heart diseases have been shown to have neurological deficits [4–6]. While various drug classes reduce mortality and morbidity in cancer patients, many of these treatments present life-threatening adverse effects, particularly cardiotoxicity and/or neurotoxicity. Drug-induced cardiotoxicity is often characterized by impairment in left ventricular function, often leading to heart failure [7]. Drug-induced neurotoxicity is often associated with debilitating neuropathic pain mediated by damage to neurons of the peripheral nervous system [8]. The onset of cardiotoxicity and neurotoxicity following cancer chemotherapy may be either acute or chronic [9–11], and consequently, clinical trials may fail to detect some toxic side effects [12, 13]. Drug-induced neurotoxicity may also involve damage to neurons caused indirectly by other cells in the body [14].

Interactions between the cardiac and neural systems have important clinical implications. The heart is actively innervated by sympathetic and parasympathetic fibers of the autonomic nervous system, and neural regulation of the cardiovascular system is well established [15, 16]. Disruption of cardiac innervation is often associated with autonomic dysregulation and neuropathy and has been shown to contribute to pathologies, including silent myocardial ischemia/infarction [17], myocardial hypertrophy, metabolic dysfunction [18], and heart failure [19]. These conditions are characterized by increased arrhythmic incidence and are associated with either enhanced or reduced sympathetic function [20–22]. Conversely, the understanding of how cardiac cells regulate neuronal cells remains limited. Existing studies have largely focused on classic trophic factors released from the heart, such as nerve growth factor (NGF). NGF has been shown to support the maturation and survival of sympathetic neurons innervating the heart through retrograde signaling from cardiomyocytes [23]. However, how cardiac metabolism contributes to neuronal health remains poorly understood.

Sympathetic nerves play crucial roles in the regulation of every heartbeat [24]. PC12, a rat pheochromocytoma cell line, can be induced to differentiate into a sympathetic neuronal phenotype by treatment with NGF and is widely used for conducting neurite outgrowth assays [25]. Some studies have also found the ability of conditioned media collected from cultures of other cell lines to induce neurodifferentiation of PC12 [26, 27]. Direct contact co-culturing of PC12 with other cell lines has also been found to induce neurodifferentiation [28]. Co-culturing PC12 with the widely used H9C2 cardiac myoblast cell line has been found to increase PC12 proliferation, under treatment with palmitic acid [29]. *In vitro* cardiac cell-based models have demonstrated the cardiotoxic effects of multiple, common Food and Drug Administration (FDA)-and Health Canada-approved, commercially available anti-cancer agents. These include, but are not limited to, anthracyclines (doxorubicin (DOX), daunorubicin, epirubicin), taxanes (paclitaxel), tyrosine kinase inhibitors (dasatinib, lapatinib, nilotinib, ponatinib, sorafenib sulfate), proteasome inhibitors (ixazomib citrate, bortezomib), protein kinase inhibitors (pemigatinib), PARP inhibitors (olaparib), etc. [30, 31]. Among these, anthracyclines represent one of the most well-established classes of cardiotoxic agents, inducing dose-dependent cardiomyopathy that can progress to heart failure [7, 31]. DOX, a classic anthracycline, has been extensively shown to induce cardiomyocyte injury through mechanisms including oxidative stress, mitochondrial dysfunction, and metabolic dysregulation [11, 30, 31].

Many cardiotoxic agents, including DOX [31], paclitaxel [30], and oxaliplatin [32], are also neurotoxic and have been found to have an inhibitory effect on neurite outgrowth of PC12 cells *in vitro* [33–35]. Since the majority of anti-cancer drug toxicity research is centered on direct damage to a particular cell type, mechanisms of secondary neurotoxicity induced by drug-injured cardiac cells remain unclear. It is known that the heart plays an important role in neuropathology [4–6]. Disruption of trophic factor signaling, including reduced NGF content in the heart following cardiotoxic injury, has been implicated in chemotherapy-associated neurotoxicity [14]. Nonetheless, trophic factor disruption may not fully account for the range of cardiomyocyte-derived signals that may impact neuronal function. Particularly, it remains unclear how changes in cardiomyocyte metabolic state after cardiotoxic injury generate aberrant signals that contribute to neuronal dysfunction.

Human-induced pluripotent stem cell-derived cardiomyocytes and sympathetic neurons (hiPSC-CMs and hiPSC-SNs) offer greater physiological and clinical relevance than traditional animal cell culture models for studying drug-induced toxicities. Existing co-culture models comprised of hiPSC-CMs and hiPSC-SNs recapitulate many aspects of sympathetic cardiac innervation, with the formation of functional neuro-cardiac junctions [36, 37] and the ability of the hiPSC-SNs to regulate the beating rate of the hiPSC-CMs [36, 37]. HiPSC-based neuro-cardiac co-culture systems could serve as a great model to investigate how cardiotoxicity impacts neurons, particularly through conditioned media-based approaches to investigate paracrine-mediated effects.

Importantly, targeted therapeutic strategies aimed at preventing or reversing cardiac-mediated secondary neurotoxicity do not currently exist. Identifying cardiomyocyte-derived metabolites that influence neuronal health could reveal novel therapeutic targets for chemotherapy-associated neurological complications.

## MATERIALS AND METHODS

### Mammalian Cell Culture

H9C2 rat cardiac myoblasts, originally obtained from ATCC,were cultured in high-glucose DMEM (Thermo Fisher Scientific) with 10% FBS (Thermo Fisher Scientific) and 1% penicillin-streptomycin (Sigma-Aldrich). The H9C2 cells were passaged regularly at 80-90% confluency using 0.25% trypsin-EDTA (Thermo Fisher Scientific). PC12 rat pheochromocytoma cells were obtained from ATCC and cultured in RPMI-1640 (Thermo Fisher Scientific), with 10% heat-inactivated horse serum (Sigma-Aldrich), 5% FBS, and 1% penicillin-streptomycin. The PC12 cells were routinely passaged every 4-6 days via centrifugation, with cell clumps dispersed by trituration with an 18-gauge needle. All cells were maintained at 37 °C and 5% CO_2._

### Cardio-Neuronal Co-culture Model

To generate a direct cardio-neuronal co-culture, PC12 cells (450,000 for 6-well, 25,000 for 96-well) were seeded in triplicate in tissue culture-treated plates coated with type 1 bovine collagen (Sigma-Aldrich) and allowed to attach overnight. On the following day, designated as PC12 day 1, H9C2 cells (80,000 for 6-well, 10,000 for 96-well) were added to the wells. The cells were co-cultured for 7 days with a media change every 48 hours. To generate an indirect co-culture, PC12 cells were treated with conditioned media (CM) collected from H9C2 cells for 7 days, with fresh CM added every 48 hours. CM was prepared by initial centrifugation for 6 minutes at 1000 x g, after which the supernatant was collected and centrifuged again for 5 minutes at 1000 x g [38]. On day 7, cells were fixed, and immunostaining for neurofilament light-chain (NF-L) was performed as described below.

### Quantification of Neurite Outgrowth

Brightfield images of co-cultures at day 7 were captured using a Zeiss Primovert inverted microscope equipped with a 20x objective, an Axiocam 5s camera, and ZEN 3.2 software (Zeiss). For neurite outgrowth quantification, 3 random fields were imaged per well [39, 40]. Images were analyzed using the NeuronJ plugin in ImageJ to obtain neurite length measurements [41–44]. Additionally, the number of neurite-bearing cells per imaging field was totaled as a complementary metric [45, 46].

### Immunofluorescence Staining

Primary antibodies were purchased from Santa Cruz Biotechnology and diluted 1:50 in 3% bovine serum albumin (BSA, GoldBio). Alexa Fluor-conjugated secondary antibodies were purchased from Thermo Fisher Scientific and diluted 1:1000 in 3% BSA. Cells were fixed with 4% paraformaldehyde (PFA) and permeabilized with 0.1% Triton X-100 (Thermo Fisher Scientific). Cells were blocked with 3% BSA and incubated with the primary antibody, followed by the corresponding fluorophore-conjugated secondary antibody. Cells were counterstained with 4′,6-diamidino-2-phenylindole (DAPI, MedChemExpress). Fluorescent images were captured using Agilent BioTek Cytation 5 and Gen5 software (Agilent). For in-cell ELISA assays, following immunostaining with the primary antibody and Alexa Fluor 594 secondary antibody, fluorescence was quantified using an Agilent BioTek Synergy HTX microplate reader (Ex/Em: 560/600 nm) in endpoint mode.

### Determination of IC50 Values of Anti-cancer Compounds on Cardiac Cells

H9C2 cells were seeded in quadruplicate in 96-well plates at a density of 10,000 cells/well and allowed to attach overnight. Twenty-four hours later, the cells were dosed with the following concentrations of DOX (MedChemExpress), ripretinib (Adooq Biosciences), mobocertinib (Adooq Biosciences), and quizartinib (Adooq Biosciences): 1 nM, 10 nM, 100 nM, 1 *μ*M, 5 *μ*M, 10 *μ*M, 25 *μ*M, and 50 *μ*M. After a 24-hour exposure period, the Cell Titer Glo (Promega) cell viability assay was performed according to the manufacturer’s instructions, and luminescence readings were obtained.

### Evaluation of Neurite Outgrowth Following Exposure to Injured Cardiac Cells

H9C2 cells were seeded in triplicate in 6-well plates at a density of 80,000 cells/well and allowed to attach overnight. Twenty-four hours later, the cells were dosed with 10 and 50 *μ*M DOX (MedChemExpress) for either 24 or 48 hours. The 10 *μ*M concentration was used as the primary experimental condition based on preliminary studies demonstrating significant effects on neurite outgrowth, as well as prior *in vitro* cardiotoxicity models [31, 47–49], while 50 *μ*M was included as a higher-severity injury condition. Three additional anti-cancer compounds, mobocertinib, ripretinib, or quizartinib (Adooq Biosciences) were applied to H9C2 cells, at concentrations selected based on their respective IC50 values: mobocertinib at 5, 10.61, and 50 *μ*M; ripretinib at 1, 12.25, and 50 *μ*M; and quizartinib at 5, 11.01, and 50 *μ*M. Following drug exposure, the cells were rinsed with 1x DPBS (Boston Bioproducts) and provided with fresh media. After an additional 24 hours of recovery, the H9C2 cells were either directly co-cultured with PC12 cells or used for CM collection.

In parallel, PC12 cells (450,000/well, in triplicate) were differentiated by treatment with conditioned media derived from healthy H9C2 cells on PC12 day 1. On PC12 day 3, cells were exposed either to injured H9C2 cells or conditioned media collected from injured H9C2 cells. The cells were co-cultured until PC12 day 7, with media changes occurring every 48 hours. Images were then captured, and neurite outgrowth quantification was performed as previously described.

### Evaluation of Neurite Outgrowth Controlling for Cardiac Cell Loss Following Injury

To ensure that reduced PC12 outgrowth observed in co-culture with drug-injured H9C2 cells was not solely caused by a decreased number of H9C2 cells, a separate cell number-matched experiment was performed with DOX, consistent with its use as a standard cardiomyocyte injury model for this study. Following treatment of H9C2 cells with 10 and 50 *μ*M DOX for 24 hours, the number of viable remaining H9C2 cells was quantified. An equal number of unexposed, healthy H9C2 cells was then seeded into co-culture with PC12 cells under the same conditions previously mentioned for direct co-cultures, and the co-culture was continued until day 7. For indirect co-culture conditions, conditioned media was collected from H9C2 cultures normalized to the same cell number and applied to PC12 cells. All groups were assayed in triplicate. Imaging and neurite quantification were performed as previously described.

### Untargeted Metabolomics

For untargeted metabolomic analysis, monoculture conditions were established in triplicate 6-well plates, and conditioned media was collected from selected DOX-treated co-culture conditions described above. The following groups were included: (1) PC12 only, (2) H9C2 only, (3) PC12 + H9C2 (direct co-culture), (4) PC12 + CM from H9C2 (indirect co-culture), (5) PC12 + injured H9C2 (10 *μ*M DOX, 24 hour incubation), (6) PC12 + CM from injured H9C2 (10 *μ*M DOX, 24 hour incubation), (7) PC12 + CM from injured H9C2 (10 *μ*M DOX, 48 hour incubation). Both monocultures and co-cultures were maintained for 7 days, and imaging and neurite quantification were completed as previously described. Following imaging, conditioned media was collected from the cells and centrifuged at 1000 x g for 1 minute at 4°C. The supernatant was collected and snap frozen in liquid nitrogen, and the samples were stored at -80 °C before shipping for analysis.

Untargeted metabolomics and analysis were performed by MtoZ Biolabs [50]. Samples were blindly processed prior to the analysis. Metabolites were extracted using methanol, with sonication and incubation at -20 °C followed by a centrifugation step. The supernatants were collected and used for further analysis. The metabolomic profiling was completed using liquid chromatography tandem mass spectrometry (LC-MS/MS) on a Thermo Fisher Scientific Q Exactive mass spectrometer with a Thermo Fisher Scientific U3000 liquid chromatography system. Chromatographic separation was performed using hydrophilic interaction liquid chromatography (HILIC).

The raw data were processed by MtoZ Biolabs using Progenesis QI software for peak alignment, noise removal, and metabolite identification and quantification. Metabolite annotation was performed using custom-built databases as well as public databases, such as the Human Metabolome Database (HMDB). Prior to statistical analysis, the data were normalized. Total value normalization and missing value imputation were performed, where missing values were replaced with one-half of the minimum non-zero value detected in the dataset. Clustering heatmap analysis was conducted using the preprocessed data. Multivariate analyses, including principal component analysis (PCA), partial least squares discriminant analysis (PLS-DA), and orthogonal partial least squares discriminant analysis (OPLS-DA), were used to assess global metabolic differences between the groups.

Differentially expressed metabolites were identified based on variable importance in projection (VIP) scores from OPLS-DA, unpaired t-tests, and fold-change analysis. P-values obtained from unpaired t-tests were adjusted for multiple comparisons using the false discovery rate (FDR) method with the Benjamini-Hochberg correction. Metabolites with VIP > 1.0, FDR-adjusted P < 0.05, and fold change ≥ 1.5 were considered upregulated, while metabolites with VIP > 1.0, FDR-adjusted P < 0.05, and fold change ≤ 0.67 were considered downregulated.

### Pharmacologic Inhibition of Metabolic Pathways

PC12 cells were seeded in triplicate in 96-well plates at a density of 25,000 cells/well and allowed to attach overnight. H9C2 cells were seeded at a density of 10,000 cells/well for the generation of CM. Candidate cardio-neuronal signaling metabolites were identified from metabolomic profiling, and associated pathways were inferred from prior literature to guide selection of pharmacologic inhibitors. Inhibitors (MedChemExpress) were applied to both PC12 and H9C2 cells in co-culture.

D-mannosamine (5 and 500 *μ*M, and 1 mM) was used to disrupt early steps in glycosylphosphatidylinositol synthesis, in order to reduce the formation of downstream intermediates including 6-(α-D-glucosaminyl)-1D-myo-inositol [51–53]. Zileuton (1, 50, and 100 *μ*M) was used a 5- lipoxygenase inhibitor to modulate arachidonic acid metabolism and downstream production of eicosanoids. The studies by Ashton et al., Rossi et al., and Patrono led us to hypothesize that Zileuton may plausibly reduce 2,3-dinor-TXB₂ levels in cultured cells [54–56]. 3Fax-Peracetyl-Neu5Ac (100 *μ*M) was used to inhibit sialyltransferase activity, to disrupt CMP-Neu5Ac-dependent sialylation and downstream metabolism of sialic acids [57–60].Inhibitor doses were based on previously published studies [61–67].

Both cell lines were dosed with inhibitors on the day following seeding. On the next day, which corresponds to PC12 day 1, PC12 cells were treated with CM from H9C2 cells. In the continuous dosing group, inhibitors were re-applied daily throughout the duration of the co-cultures, while in the single-dose group, no additional inhibitor dose was given after the initial exposure. Vehicle-treated co-cultures were included as the controls. Additionally, a co-culture group dosed with 10 *μ*M DOX, as previously described, was included as a negative control for reduced neurite outgrowth. The co-cultures were maintained until PC12 day 7 as previously described for indirect co-culture experiments. Imaging and neurite outgrowth quantification were also conducted as previously described.

### hiPSC Culture

Female human induced pluripotent stem cells (hiPSCs) were obtained from Stanford University and cultured in StemMACS (Miltenyi Biotech) on Matrigel-coated (Corning) 6-well plates. The hiPSCs were passaged regularly at 80-85% confluency using a gentle cell dissociation reagent (GCDR, Stem Cell Technologies) at a 1:6 splitting ratio. The cells were maintained at 37° C and 5% CO_2_.

### hiPSC Differentiation into Cardiomyocytes and Sympathetic Neurons

Beating cardiomyocyte differentiation was carried out using the BMEM protein-free cardiomyocyte differentiation kit (BMEM Bio) according to the manufacturer’s instructions. Differentiation was validated by immunostaining as previously described for cardiac troponin T (cTnT) [68].

Sympathetic neuronal differentiation was performed according to previously published methods by Wu et al [69]. Briefly, hiPSCs were dissociated using GCDR and seeded at a density of 125,000 cells/ cm^2^ on Matrigel-coated 6-well plates (day 0). The cells were seeded in day 0-1 media: Essential 6 (E6) medium (Thermo Fisher Scientific) with 10 *μ*M SB431542 (MedChemExpress), 1 ng/ml BMP4 (PeproTech), 300 nM CHIR99021 (SelleckChem), and 10 *μ*M Y27632 (Enzo Life Sciences). On the following day (day 1), the media was refreshed, and Y27632 was excluded. On days 2-10, the cells were cultured in E6 medium containing 10 *μ*M SB and 0.75 *μ*M CHIR99021, with a media change occurring every other day.

On day 10, cells were washed with 1X DPBS and dissociated by incubating with Accutase (Innovative Cell Technologies) for 20 minutes at 37 °C and 5% CO_2_. The dissociated cells were pooled into a 50 ml conical tube, which was topped up with DPBS. Cells were then centrifuged at 200 x g for 4 minutes at room temperature, followed by resuspension in day 10-14 media: Neurobasal medium (Thermo Fisher Scientific) with B27 supplement without vitamin A (Thermo Fisher Scientific), N2 supplement, (Thermo Fisher Scientific), 2 mM L-glutamate (Thermo Fisher Scientific), 3 *μ*M CHIR99021, and 10 ng/ml FGF2 (MedChemExpress). Spheroids were formed by plating 0.5 × 10^6 cells per well in 24-well ultra-low attachment plates (SPL Life Sciences).

Spheroids were fed every other day until day 15, at which point they were replated for final sympathetic neuronal differentiation. Twenty-four-well plates were coated with poly-D-lysine, laminin, and fibronectin (all from Thermo Fisher Scientific). Spheroids were split 1:2 and plated on the coated plates in sympathetic neuronal maturation medium consisting of Neurobasal medium with B27 supplement with vitamin A (Thermo Fisher Scientific), N2 supplement, 2 mM L-glutamate, 25 ng/ml GDNF (MedChemExpress), 25 ng/ml BDNF (MedChemExpress), 25 ng/ml NGF (MedChemExpress), 200 *μ*M ascorbic acid (Fujifilm Wako Chemicals), 0.2 mM dbcAMP (MedChemExpress). Neurons were fed every other day until day 20, from which point they were fed weekly by performing a partial media change. Differentiation was validated by immunostaining as previously described for tyrosine hydroxylase (TOH) [70].

### hiPSC Model of Cardiac-induced Neuronal Dysfunction

To generate a hiPSC model of cardiac-induced neuronal dysfunction, hiPSC-CMs and hiPSC-SNs were differentiated as described above. Day 20 hiPSC-CMs were treated with 250 nM and 1 *μ*M DOX for 24 hours, doses selected according to prior hiPSC-based cardiotoxicity studies [71, 72]. Following the 24-hour incubation, the hiPSC-CMs were washed and incubated in fresh media for 24 hours, after which conditioned media was collected as previously described. Conditioned media was applied to day 26 hiPSC-SNs at a 1:1 ratio with sympathetic neuronal maturation media. All conditions were performed in triplicate. After 48 hours of exposure to conditioned media from the hiPSC-CMs, hiPSC-SN media was collected for norepinephrine (NE; noradrenaline) quantification, and cells were fixed for immunostaining.

### βIII-tubulin Immunofluorescence and Image Analysis

Fixed hiPSC-SNs were stained for βIII-tubulin as previously described. Fluorescent images of βIII-tubulin were acquired from six random fields per well with uniform imaging settings. βIII-tubulin fluorescence intensity was analyzed using ImageJ and reported as the integrated density values [73, 74].

### Norepinephrine (NE) Quantification

Measurement of NE levels in conditioned media was performed using the Invitrogen noradrenaline competitive ELISA kit (Thermo Fisher Scientific) according to the manufacturer’s instructions. Media samples were spun down at 1000 x g for 20 minutes at 4° C to remove cells and debris, and stored at -80° C prior to the analysis. Absorbance was measured and corrected by subtracting readings from media-only control samples [75].

### Statistical Analysis

Data analysis was performed using GraphPad Prism (version 10) software. Grouped comparisons were analyzed using one-way analysis of variance (ANOVA) with Tukey’s multiple comparison test, Dunnett’s test, or t-tests. P <0.05 was considered statistically significant. For the determination of IC50 values, non-linear regression analysis was performed by log-transforming concentrations and fitting the data using a four-parameter variable slope dose-response model. NE ELISA concentrations were determined by interpolation from standard curves generated using four-parameter logistic nonlinear regression analysis. Analysis for untargeted metabolomics is described above.

## RESULTS

### Co-cultured Cardiac Cells Induced Outgrowth of NF-L-positive Neurites

Brightfield images showed the presence of neurites on PC12 cells following seven days of co-culture with H9C2 cells (Fig. 1C) or conditioned media from H9C2 cells (Fig. 1D). Neurites were not observed in PC12 cultured alone (Fig. 1A). Statistical analysis confirmed a significant increase in both neurite length (Fig. 1E) and number of neurite-bearing PC12 cells (Fig. 1F) in both direct (PC12+H9C2) and indirect (PC12 + conditioned media [CM] from H9C2) co-cultures, compared to PC12 cultured alone. PC12 neurites within direct and indirect co-cultures expressed neuronal differentiation marker, NF-L [76–78] as assessed by immunofluorescence (Fig. 1). These findings confirm the ability of H9C2 cells to induce neurodifferentiation of PC12 without external trophic support.

**Figure 1:**
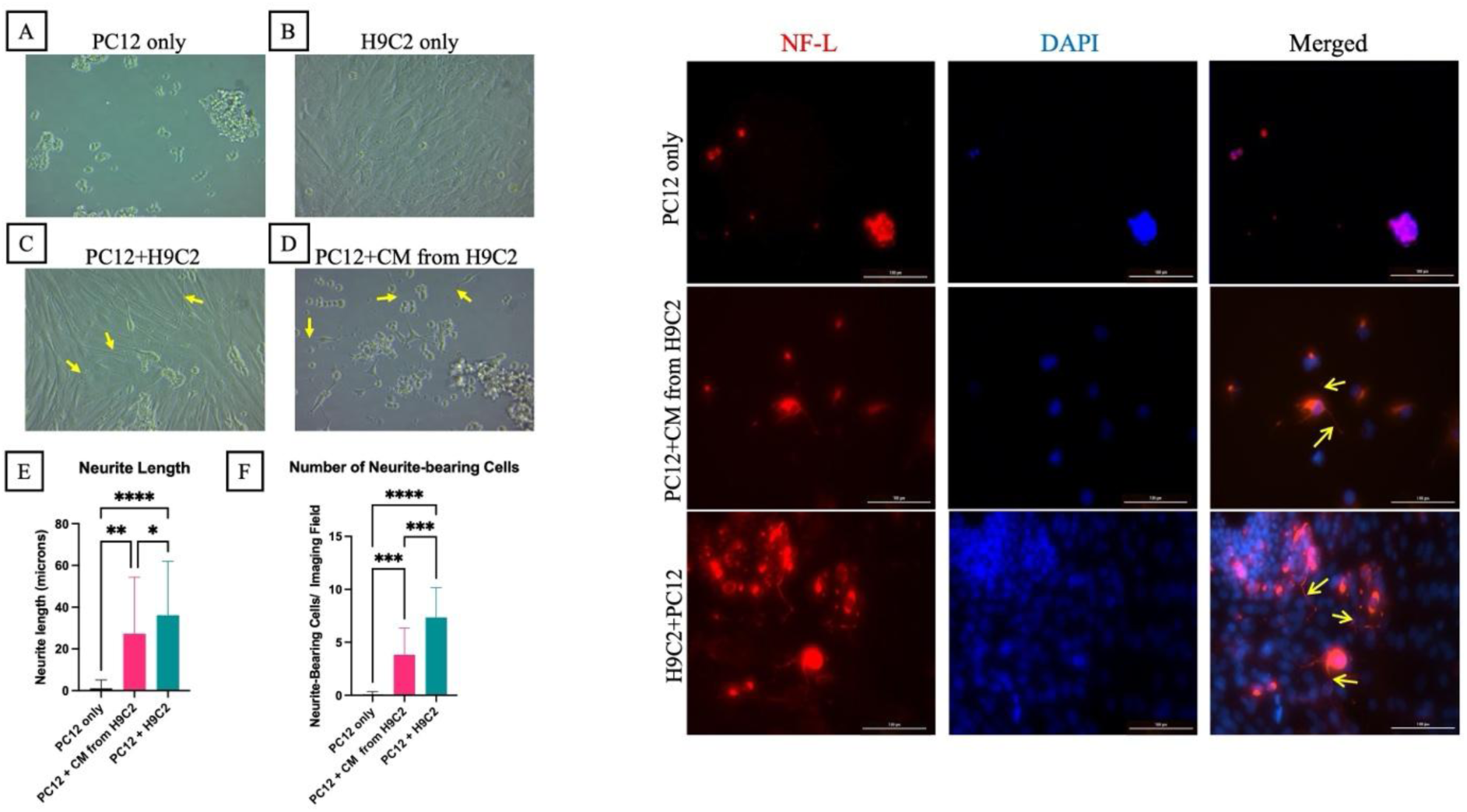
Cardiac Cells Induce Neurite Outgrowth Through Direct Contact or Conditioned Media. Left: brightfield images, captured at 20x magnification, showing **(A)** PC12 only, **(B)** H9C2 only, **(C)** PC12+H9C2 (direct co-culture), **(D)** PC12+ conditioned media (CM) from H9C2 (indirect co-culture). Arrows indicate representative neurite outgrowths. **(E)** Neurite length and **(F)** number of neurite-bearing cells significantly increased in PC12 cells co-cultured with H9C2 cells, both via direct and indirect co-culture methods, compared to PC12-only controls. Data is represented as means ± standard deviation. Data was analyzed via one-way ANOVA and Tukey’s post hoc test with GraphPad Prism. [*] p < 0.05, [**] p < 0.01, [***] p < 0.001, [****] p< 0.0001. Right: immunofluorescent staining of PC12 only, PC12+CM from H9C2, and H9C2+PC12 with neurofilament light chain (NF-L, red) and DAPI (nuclear stain, blue). Arrows indicate representative neurite outgrowths. Scale bar = 100 microns; images were captured at 10x magnification.

### DOX Administration to Cardiac Cells Reduced Neurite Outgrowth

Injuring H9C2 cells with DOX at select doses prior to co-culture with PC12 significantly decreased neurite length and the number of neurite-bearing PC12 cells compared to control co-cultures (Fig. 2). In the direct co-cultures, DOX exposure at 10 and 50 *μ*M for 24 or 48 hours on H9C2 cells significantly decreased neurite lengths (Fig. 2A and 2C) and number of neurite-bearing PC12 cells (Fig. 2B and 2D). Similarly, exposing H9C2 to DOX for 24 or 48 hours prior to co-culture with PC12 significantly reduced neurite length (Fig. 2E and 2G) and number of neurite-bearing cells (Fig. 2F and 2H) at doses of 10 and 50 *μ*M in indirect co-cultures. Representative brightfield and immunofluorescence images (Suppl. Fig. 2) support these findings, showing TOH-positive neurite outgrowth in healthy indirect co-cultures and weaker TOH expression with no neurite formation following 10 *μ*M DOX treatment. Taken together, these results demonstrate that induction of cardiotoxicity in H9C2 (either with cells or media alone) can attenuate their ability to differentiate PC12 cells.

**Figure 2:**
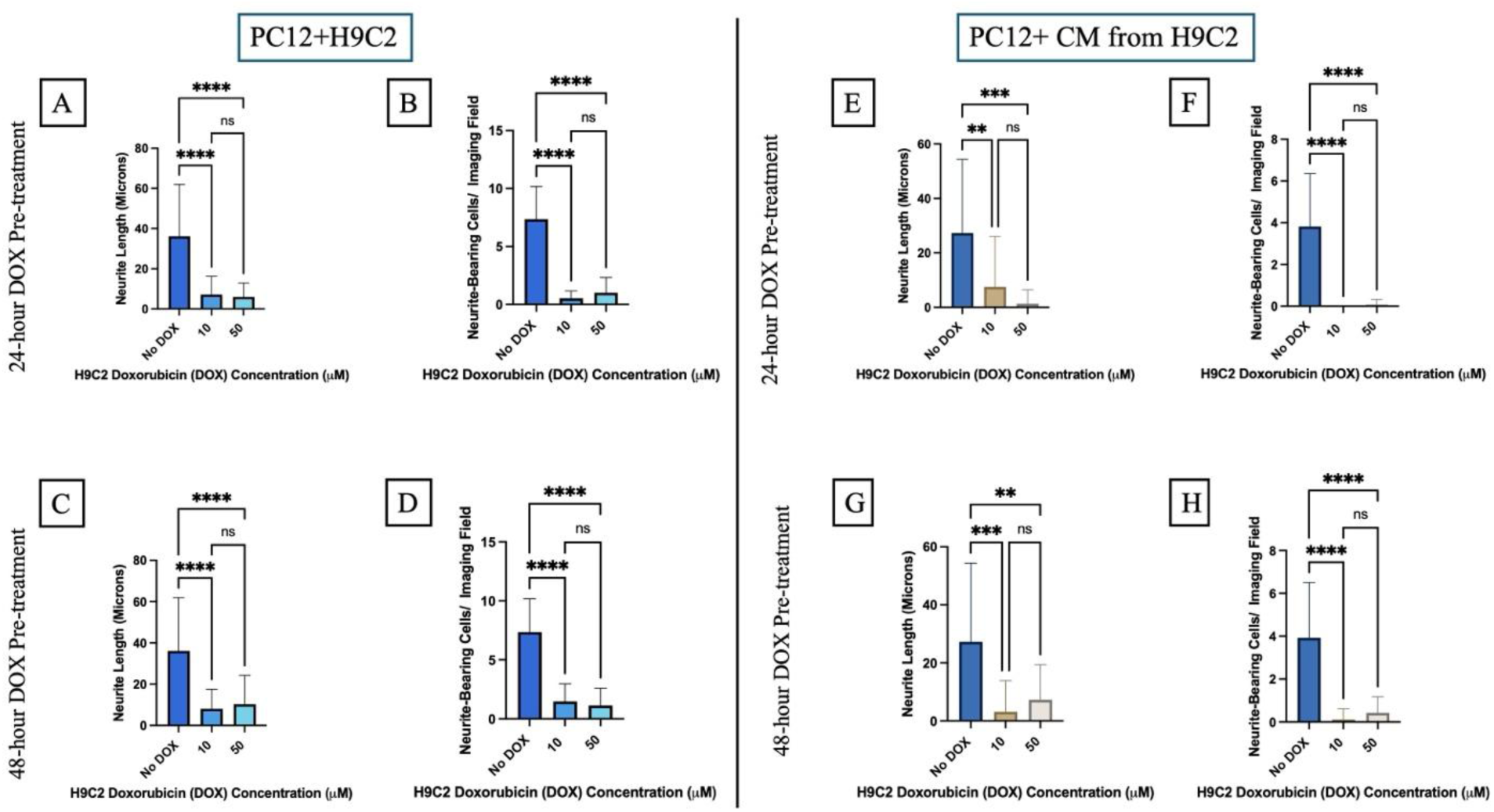
DOX Administration Inhibits Cardiac-Cell-Induced Neurite Outgrowth. Pre-treatment of H9C2 with doxorubicin (DOX) at 10 μM μM, for **(A, B, E, F)** 24 hours and **(C, D, G, H)** 48 hours significantly decreased PC12 neurite length and number of neurite-bearing PC12 in both direct co-cultures (PC12+H9C2, **A-D**) and indirect co-cultures (PC12 + conditioned media [CM] from H9C2, **E-H).** Data is represented as means ± standard deviation. Data was analyzed via one-way ANOVA and Tukey’s post hoc test with GraphPad Prism. [*] p < 0.05, [**] p < 0.01, [***] p < 0.001, [****] p < 0.0001

### Reduction in Neurite Outgrowth Following Cardiotoxic Injury Is Not Due to Cardiac Cell Loss

To verify if the reduction in neurite outgrowth observed in DOX-exposed co-cultures was not attributable to a reduced number of H9C2 cells following injury, we performed co-culture experiments controlling for DOX-induced H9C2 cell loss. Directly co-culturing PC12 cells with equivalent numbers of either healthy H9C2 cells or the number of remaining H9C2 cells following DOX injury at 10 *μ*M (Fig. 3A and 3B) or 50 *μ*M (Fig. 3C and 3D) resulted in no significant differences in neurite length (Fig. 3A and 3C) and number of neurite-bearing cells (Fig. 3B and 3D) compared to control co-cultures containing the regular number of H9C2 cells.

**Figure 3:**
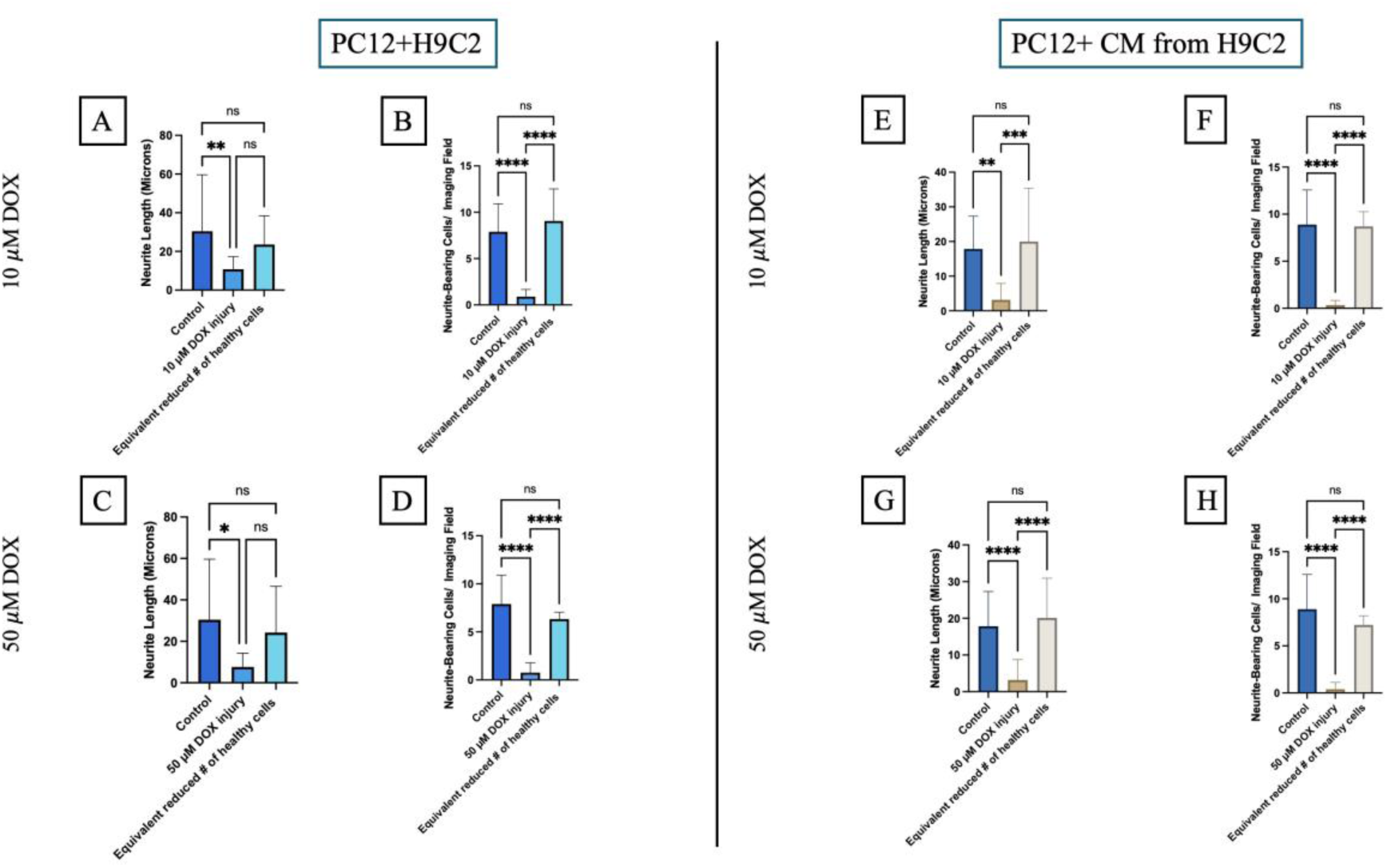
Reduced Neurite Outgrowth Following DOX Injury Is Not Due to Cardiac Cell Loss. Seeding healthy H9C2 cells at numbers equivalent to those remaining after doxorubicin (DOX) treatment for 24 hours [10 μM **(A-B)** and 50 μM **(C-D)**] did not significantly reduce (**A & C)** PC12 neurite length or **(B & D)** number of neurite-bearing PC12 cells compared to control direct co-cultures (PC12+H9C2). Similarly, in indirect co-cultures with PC12 + conditioned media [CM] from H9C2 exposed to DOX for 24 hours [10 μM **(E-F)** and 50 μM **(G-H)**], no significant reduction in neurite length **(E & G)** and number of neurite-bearing cells **(F & H)** was seen in groups with an equivalent reduced number of healthy H9C2 cells as DOX injury groups. Data is represented as means ± standard deviation (n=3). Data was analyzed using one-way ANOVA with Tukey’s post hoc test in GraphPad Prism. [*] p < 0.05, [**] p < 0.01, [***] p < 0.001, [****] p < 0.0001

Consistent with these findings, indirect co-culture experiments in which equivalent numbers of healthy H9C2 cells were similarly used showed no significant reduction in neurite length (Fig. 3E and 3G) or the number of neurite-bearing cells (Fig. 3F and 3H) compared to controls. These findings demonstrate that the reduction in PC12 outgrowth seen in co-cultures containing DOX-exposed H9C2 is not attributable to decreased H9C2 cell number, but rather to altered signaling from H9C2 cells following injury.

### Co-culture and DOX Exposure Altered Metabolite Signatures

We next examined metabolite alterations in conditioned media under direct and indirect co-culture conditions, ± DOX administration to H9C2 cells. Co-cultures used for metabolomic media collection showed reduced neurite length and number of neurite-bearing cells under DOX-exposed conditions compared to healthy controls (Fig. 2). Untargeted metabolomic profiling identified 474 metabolites across all groups (Fig. 4). Comparisons between DOX-exposed groups revealed minimal metabolic differences between direct (24-hour exposure) and indirect (both 24- and 48-hour exposures) co-culture conditions. No significant differences were observed between indirect DOX-exposed conditions at 24 and 48 hours.

**Figure 4:**
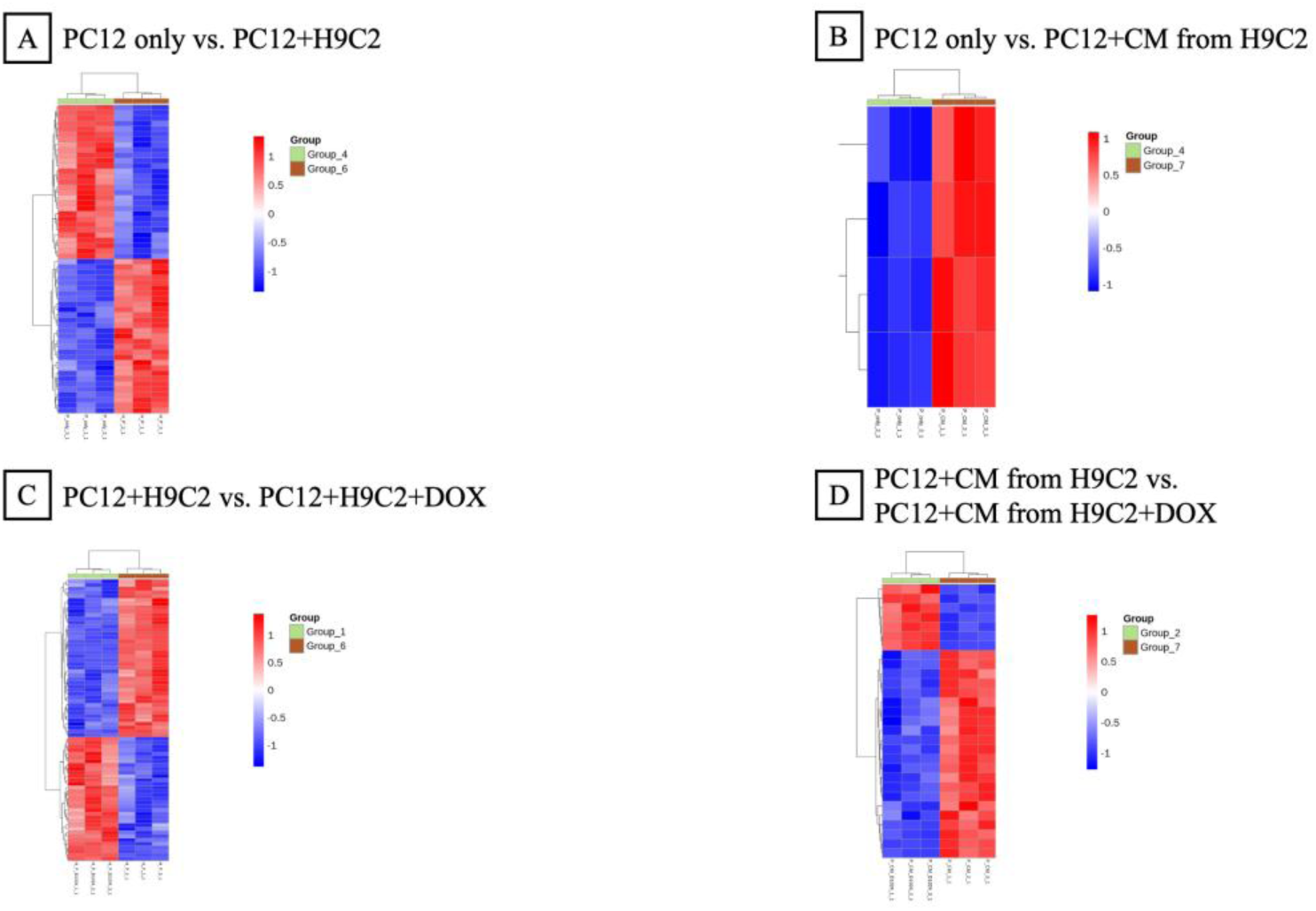
Metabolomic Profiling Reveals Key Metabolites Following DOX Injury. Heatmaps showing **(A)** changes in metabolite abundances in PC12 only monocultures compared to PC12+H9C2 (direct co-culture), **(B)** PC12 only monocultures compared to PC12+conditioned media (CM) from H9C2 cells (indirect co-culture), **(C)** PC12+H9C2 (direct co-culture) compared to PC12+H9C2 treated with 10 μM doxorubicin (DOX) for 24 hours, and **(D)** PC12 + CM from H9C2 (indirect co-culture) compared to PC12 + CM from H9C2 treated with 10 μM doxorubicin for 24 hours. Data was normalized and imputed as described in Methods, followed by hierarchical clustering. The color scale represents relative metabolite levels after scaling, with red indicating higher abundance and blue indicating lower abundance.

In contrast, larger differences in metabolic profile were observed between co-culture conditions and PC12 monocultures. Direct co-culture (PC12 + H9C2) showed 58 significantly altered metabolites compared to PC12 only cultures, while indirect co-culture (PC12 + CM from H9C2) showed 4 significant changes relative to PC12 only cultures. Notably, among these four metabolites, three were similarly altered in direct co-cultures, indicating a shared metabolic signature across both co-culture conditions.

DOX exposure increased the number of altered metabolites across comparisons. Direct DOX-exposed co-cultures showed 19 significantly altered metabolites relative to PC12 only cultures (10 upregulated, 9 downregulated), while indirect DOX-exposed co-cultures showed 3 significant alterations with 24-hour-exposure (1 upregulated, 2 downregulated) and 17 significant changes with 48-hour-exposure (10 upregulated, 7 downregulated) compared to the PC12 only group. Comparisons to unexposed co-culture conditions revealed greater changes, with up to 90 significantly altered metabolites observed. Four significantly downregulated metabolites were detected between PC12 only and H9C2 only conditions.

To identify candidate mediators of cardiac-induced neurite outgrowth, we focused on metabolites that were increased in co-culture relative to PC12 monocultures and decreased following DOX exposure on H9C2 cells, selecting these for downstream experiments. In direct co-cultures, 6-(α-D-glucosaminyl)-1-phosphatidyl-1D-myo-inositol (FC = 0.298, P = 0.031) and CMP-N-glycolylneuraminate (FC = 0.093, P = 0.036) were higher in direct co-culture than in PC12 monocultures. In indirect co-culture, CMP-N-glycolylneuraminate was also significantly elevated relative to PC12 alone (FC = 0.191, P = 0.039), suggesting that this is a common target for both co-culture methods.

Interestingly, DOX exposure on H9C2 cells prior to direct co-culture led to reduced levels of the two previous metabolites that were elevated in co-culture, 6-(α-D-glucosaminyl)-1-phosphatidyl-1D-myo-inositol (FC = 0.00005) and CMP-N-glycolylneuraminate (FC = 0.33), as well as a decrease in 2,3-dinor-TXB2 (FC = 0.32), relative to unexposed direct co-cultures. These results indicate that H9C2 DOX exposure alters the metabolic environment, affecting both previously elevated glycolylneuraminate- and phosphatidyl myoinositol-related metabolites and additional metabolites such as 2,3-dinor-TXB2.

### Pharmacologic Inhibition of Selected Metabolic Pathways Reduced Cardiac Cell-Induced Neurite Outgrowth

Inhibition of the 6-(alpha-D-glucosaminyl)-1-phosphatidyl-1D-myo-inositol pathway using D-mannosamine significantly reduced neurite length (Fig. 5A) at doses of 5 *μ*M and 500 *μ*M in indirect co-cultures receiving both a single, initial inhibitor dose as well as re-dosed co-cultures, compared to healthy controls. At a dose of 1 mM, D-mannosamine reduced neurite length only in the re-dosed group (Fig. 5A). Reduction in the number of neurite-bearing PC12 cells was observed within the re-dosed groups at 5 *μ*M, 500 *μ*M, and 1 mM (Fig. 5B), while single-dosed groups showed this reduction only at 1 mM (Fig. 5B).

**Figure 5:**
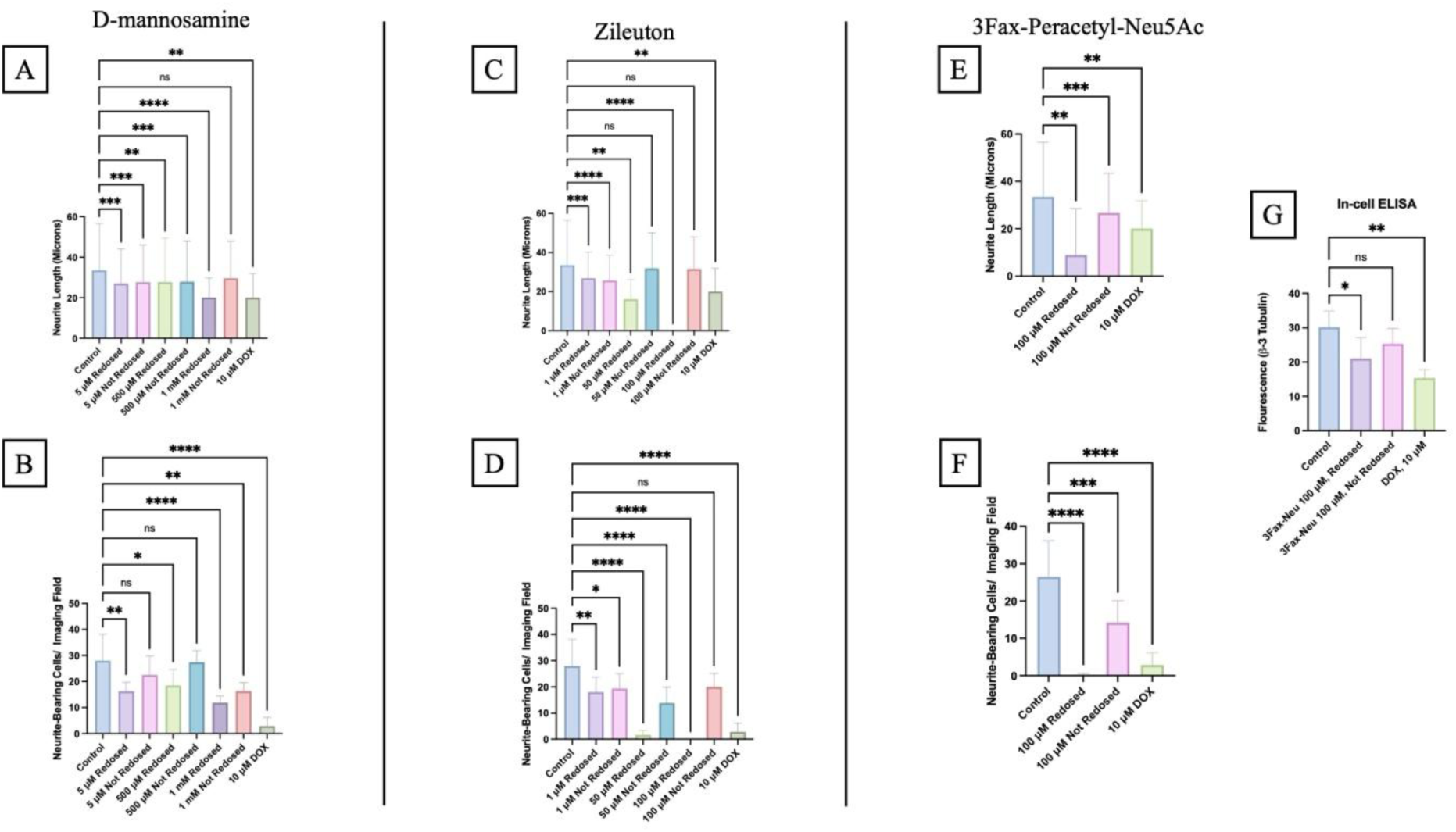
Inhibition of Select Metabolic Pathways Reduces Neurite Outgrowth. Pharmacologic inhibition of select metabolic pathways using **(A, B)** D-mannosamine, **(C, D)** Zileuton, and **(E, F)** 3Fax-Peracetyl-Neu5Ac at select concentrations significantly reduced PC12 neurite length and number of neurite-bearing cells in indirect co-cultures compared to untreated controls, either by a singular inhibitor dose or by a re-dosing regimen, as described in Methods. **(G)** In-cell ELISA showed reduced expression of βIII-tubulin in re-dosed co-cultures treated with 3Fax-Peracetyl-Neu5Ac compared to untreated controls. A co-culture group treated with 10 *μ*M doxorubicin (DOX) for 24 hours is included as a negative control for reduced neurite outgrowth. Data is represented as means ± standard deviation (n=3). Data was analyzed using one-way ANOVA with Tukey’s or Dunnett’s post hoc test in GraphPad Prism. [*] p < 0.05, [**] p < 0.01, [***] p < 0.001, [****] p < 0.0001

Similarly, inhibition of the 2,3-dinor-TXB2 pathway using Zileuton significantly reduced neurite length in all re-dosed groups (1, 50, and 100 *μ*M; Fig. 5C). In contrast, only the 1 *μ*M dose reduced neurite length in single-dosed cultures (Fig. 5C). Reduction in the number of neurite- bearing cells was observed across all doses and dosing regimens except for the single 100 *μ*M condition (Fig. 5D).

Inhibition of the CMP-N-glycolylneuraminate pathway (indirectly) using 100 *μ*M 3Fax-Peracetyl-Neu5Ac reduced neurite length (Fig. 5E) and number of neurite-bearing cells (Fig. 5F) in both single-dosed and re-dosed co-cultures. Consistent with these morphological changes, βIII-tubulin levels were significantly decreased in the re-dosed group as measured by in-cell ELISA (Fig. 5G).

As expected, DOX (10 *μ*M) exposure on H9C2 for 24 hours consistently reduced neurite length, the number of neurite-bearing cells, and βIII-tubulin levels (Fig. 5), confirming its role as a negative control. Together, these results indicate that multiple metabolic pathways support cardiac-induced neurite outgrowth, and their disruption impairs it.

### Recent Approved Anti-Cancer Agents Attenuate Cardiac-Cell-Induced Neurite Outgrowth

Recently FDA- and Health Canada-approved anti-cancer therapies (2020 or later), including the type II KIT/PDGFRA switch-control tyrosine kinase inhibitor ripretinib, the epidermal growth factor receptor (EGFR) exon 20 insertion tyrosine kinase inhibitor mobocertinib, and the FLT3-selective type II tyrosine kinase inhibitor quizartinib, have emerged as cardiotoxic agents associated with hypertension, heart failure, myocardial infarction, QT prolongation, and atrial fibrillation [79–85]. However, the understanding of their role in cardiac neurotoxicity is limited. We dosed H9C2 cells in indirect co-cultures with ripretinib (Fig. 6A-B), mobocertinib (Fig. 6C-D), and quizartinib (Fig. 6E-F) for 24 hours. All three of these agents reduced H9C2 cell viability in a dose-dependent manner (Suppl. Fig. 1A).

**Figure 6:**
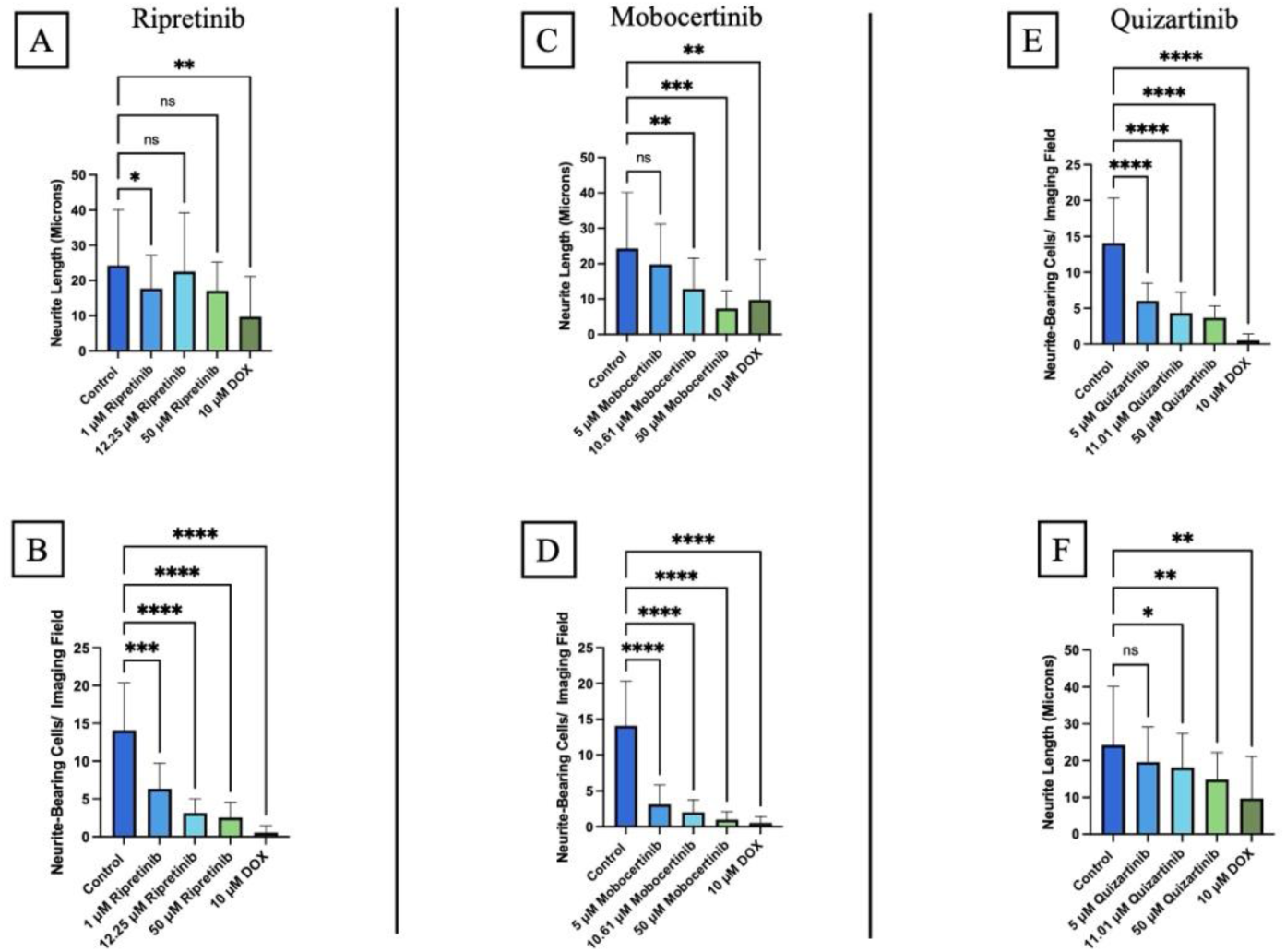
Recent Anti-cancer Agents Attenuate Cardiac-Cell-Induced Neurite Outgrowth. Pre-treatment of H9C2 for 24 hours with ripretinib **(A, B)**, mobocertinib **(C, D),** and quizartinib **(E, F)** significantly decreased PC12 neurite length **(A, C, E)** and number of neurite-bearing PC12 **(B, D, F)** in indirect co-cultures, compared to DOX-free controls, at select doses. A co-culture group treated with 10 *μ*M doxorubicin (DOX) for 24 hours is included as a negative control for reduced neurite outgrowth. Data is represented as means ± standard deviation (n=3). Data was analyzed via One-Way ANOVA and Tukey’s post hoc test with GraphPad Prism. [*] p< 0.05, [**] p < 0.01, [***] p < 0.001, [****] p < 0.0001

Ripretinib significantly reduced neurite length at a dose of 1 *μ*M (Fig. 6A) and reduced the number of neurite-bearing cells at all three doses (1, 12.25, and 50 *μ*M; IC50 = 12.25, Fig. 6B and Suppl. Fig. 1B). Mobocertinib reduced neurite length at doses of 10.61 and 50 *μ*M (Fig. 6C) and also reduced the number of neurite-bearing cells at all doses (1, 10.61, and 50 *μ*M; IC50 = 10.61, Fig. 6D and Suppl. Fig. 1B). Quizartinib reduced neurite length at doses of 11.01 and 50 *μ*M (Fig. 6E) and reduced the number of neurite-bearing cells at all doses (1, 11.01, and 50 *μ*M; IC50 = 11.01, Fig. 6F and Suppl. Fig. 1B).

As expected, DOX (10 *μ*M; IC50=1.08 *μ*M, Fig. 6 and Suppl. Fig. 1B) significantly reduced both neurite outgrowth and the number of neurite-bearing cells, consistent with its role as a negative control. Together, these results indicate that several targeted cancer therapies can impair cardiac-induced neurite outgrowth in this co-culture system, supporting the potential applicability of this model for identifying cardiotoxic agents that may indirectly induce neurotoxicity.

### DOX-induced Cardiotoxicity Impairs Sympathetic Neuronal Structure and Function in hiPSC Model

To extend our model of cardiac-induced neurotoxicity to human cells, we employed a conditioned media-based co-culture of hiPSC-SNs and DOX-exposed iPSC-CMs (Fig. 7). Differentiation of hiPSC-CMs and hiPSC-SNs was validated by cTnT and TOH immunostaining, respectively (Suppl. Fig. 3). Conditioned media from hiPSC-CMs treated with 250 nM or 1 μM DOX for 24 hours, when applied to hiPSC-SNs, significantly reduced βIII-tubulin fluorescence intensity (Fig. 7D) and NE secretion (Fig. 7E) in hiPSC-SNs compared to controls. Collectively, these results demonstrate that conditioned media from DOX-exposed hiPSC- CMs impairs both structural and functional properties in hiPSC-SNs.

**Figure 7:**
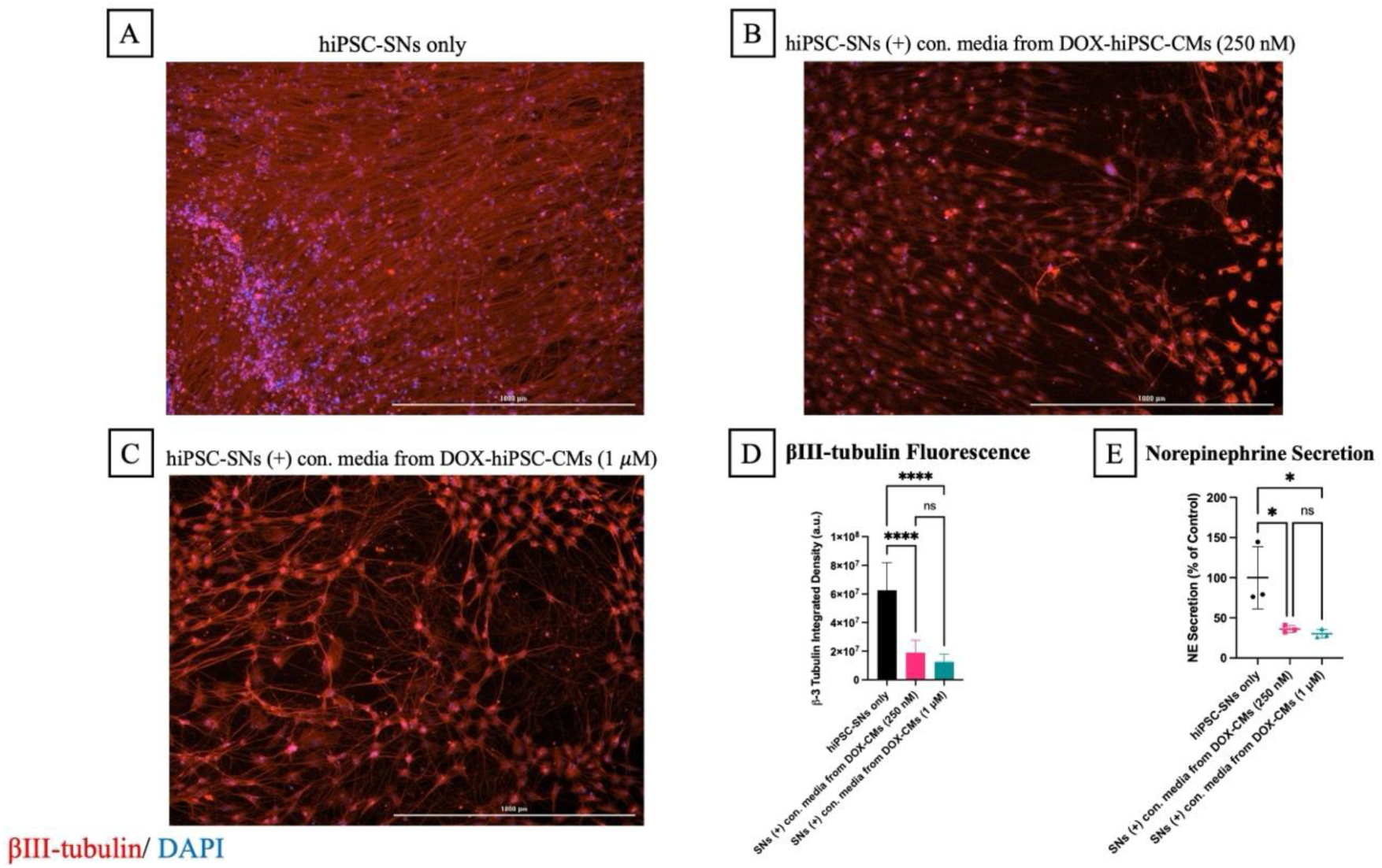
DOX-induced Cardiotoxicity Alters Sympathetic Neurons in hiPSC Model. Representative images of hiPSC sympathetic neurons (SNs) cultured alone **(A)**, or **(B)** treated with conditioned media from hiPSC-cardiomyocytes (CMs) exposed to doxorubicin (DOX) at 250 nM **(C)** or 1 μM **(D)** for 24 hours. Cells were stained for βIII-tubulin (red) and DAPI (nuclear stain, blue). Scale bar = 1000 microns; images were captured at 4x magnification. **(D)** Fluorescence intensity of βIII-tubulin and **(E)** norepinephrine secretion significantly decreased in hiPSC-SNs treated with media from DOX-exposed CMs compared to hiPSC-SNs cultured alone. Data is represented as means ± standard deviation (n=3). Data was analyzed via One-Way ANOVA and Tukey’s post hoc test with GraphPad Prism. NE ELISA data is presented as percent of untreated controls. [*] p < 0.05, [**] p < 0.01, [***] p < 0.001, [****] p < 0.0001

## DISCUSSION

### Cardiac Cells Promote Neurite Outgrowth Directly or Indirectly

Many anti-cancer therapies are associated with both cardiotoxic and neurotoxic effects, yet the contribution of cardiotoxic injury to neuronal dysfunction remains poorly understood. To date, evidence for indirect cardiac neurotoxicity has largely focused on reduced cardiac production of NGF following cardiotoxic injury [14]. In the present study, we established an *in vitro* cardio-neuronal co-culture model and demonstrated that cardiac cells promote neurite outgrowth, and cardiomyocyte injury significantly impairs this outgrowth process. Our findings further suggest that alterations in cardiac metabolic signaling may contribute to this disruption, identifying a potential non-canonical mechanism linking cardiotoxic injury to neurotoxicity. In our co-culture model, cardiac cells induced neurite outgrowth in the absence of exogenous trophic support, indicating that cardiomyocytes provide signals capable of supporting neuronal differentiation and outgrowth. Prior studies on cardio-neuronal communication have largely focused on canonical trophic factors, which are important for the maintenance of cardiac innervation and neuronal survival. NGF transfer has been shown to occur by direct cellular contact, through retrograde transport across the neurocardiac junction via TrkA receptors [23]. Other neurotrophins include glial cell line-derived neurotrophic factor (GDNF), which regulates neuronal growth, axon guidance, and cardiac innervation [86, 87], and brain-derived neurotrophic factor (BDNF), which plays a role in neuronal growth and cardiac remodeling following injury [87, 88]. Notably, our findings in both direct and indirect co-culture systems suggest that cardiac-derived signals extend beyond these classic neurotrophins, with indirect co-cultures highlighting a paracrine component of cardio-neuronal signaling.

### Anticancer Cardiotoxic Injury Impairs Cardiac-induced Neurodifferentiation

DOX Cardiotoxic injury of H9C2 cells impaired their ability to induce neurite outgrowth in PC12 cells, indicating that cardiomyocyte health directly influences neuronal differentiation in our co-culture system. PC12 cells provide a well-established *in vitro* model of sympathetic neuronal differentiation, in which neurite outgrowth reflects key processes that underlie neuronal integrity, axon growth, and nerve regeneration [89–92]. Neurotoxicity associated with DOX has been well documented in vivo, resulting in cognitive impairment and neuronal damage despite its limited penetration of the blood–brain barrier [93–95]. These findings have largely been interpreted as reflecting direct effects on neuronal cells, including oxidative stress, mitochondrial dysfunction, and neuroinflammatory signaling [94, 96, 97]. Accordingly, *in vitro* studies employing PC12 cells to study neurotoxicity have demonstrated that direct exposure of neuronal cells to chemotherapeutic agents, including DOX, inhibits neurite outgrowth [33–35]. Notably, the neurotoxic effects observed in our model were also seen in indirect co-cultures in which PC12 cells were not directly exposed to DOX. This suggests that paracrine cardiac signals may play crucial roles in cardiac-induced neurotoxicity by potentially altering the composition of the cardiac secretome.

To determine whether cardiac-induced neurotoxicity extends beyond DOX, we evaluated the effects of three more recently (2020 or later) FDA and Health Canada-approved anti-cancer drugs in our co-culture system. At a series of dosages tested, administration of ripretinib, mobocertinib, and quizartinib to H9C2 cells significantly reduced neurite outgrowth. All three drugs are tyrosine kinase inhibitors with documented cardiac adverse effects [79–85], yet are not classically characterized by direct neurotoxic effects. Although mobocertinib was later voluntarily withdrawn after a phase-III trial failed to meet its primary efficacy endpoint [85], no new safety concerns were reported for the drug. However, our results suggest that mobocertinib may contribute to cardiac-induced neurotoxicity. Collectively, our findings suggest that cardiomyocyte-mediated mechanisms may be underappreciated as contributors to neurotoxicity. In addition, further investigating the neurotoxicity profile of these three newer anti-cancer drugs will be important in understanding any possible long-term neurotoxic effects for currently approved indications, or if the drugs are considered for newer indications not approved so far.

### Metabolic Pathways Implicated in Cardiac-induced Neurite Outgrowth

Untargeted metabolomic profiling of conditioned media revealed unique metabolic signatures in direct and indirect co-cultures compared to PC12 monocultures, as well as across DOX-exposed co-cultures compared to healthy controls. In healthy indirect co-cultures, four metabolites were upregulated compared to PC12 monocultures, three of which were also upregulated in direct co-cultures relative to PC12 monocultures. This finding suggests that similar cardiomyocyte-derived metabolites may be critical for neuronal differentiation processes across both co-culture conditions. However, the relative contribution of direct cell-cell interactions versus secreted metabolic factors to neurite outgrowth remains unclear and warrants further investigation. CMP-N-glycolylneuraminate is an activated sialic acid, a class of molecules added to cell surface lipids and proteins known for modulating cellular interactions with the extracellular environment. Sialylation of cell surface molecules, particularly polysialylation, has been shown to regulate neuronal interactions and support neurite extension [98–100], yet the specific role of CMP-linked sialic acids in neurite extension has not been well defined. In this context, the increased levels of CMP-N-glycolylneuraminate observed in both direct and indirect co-culture conditions suggest that cardiac cells may promote a more growth-supportive environment, at least in part, through pathways related to sialic acid metabolism. Importantly, pharmacological targeting of this pathway was sufficient to reduce neurite outgrowth, supporting a possible functional role for this class of metabolites in mediating cardiac-induced neuronal outgrowth.

6-(α-D-glucosaminyl)-1-phosphatidyl-1D-myo-inositol is a glycosylated phosphatidylinositol intermediate. Phosphatidylinositol derivatives are important regulators of signaling cascades that exert control over cytoskeletal dynamics and membrane remodeling [101–103], which are processes essential for successful neurite extension and growth cone progression [104, 105]. In our model, this metabolite was increased in direct co-culture compared to PC12 monoculture and reduced following DOX exposure. This finding is suggestive that cardiac phosphatidylinositol signaling may, in part, have a role in supporting neuronal growth. Although phosphatidylinositol species are membrane-associated, their presence in conditioned media likely reflects membrane turnover or release through extracellular vesicles, which contain lipid components derived from the donor cell [106]. Notably, our study shows that inhibition of phosphatidylinositol-associated pathways resulted in reduced neurite length and neurite-bearing cells, supporting a functional role for lipid-derived signaling in cardiac-induced neurite outgrowth. 2,3-dinor-TXB2 is a metabolite of thromboxane A2, a lipid derived from arachidonic acid metabolism. It is commonly used as an indicator of thromboxane production [107]. Thromboxane A2 belongs to the broader class of eicosanoids, which are lipid mediators known to regulate diverse cellular signaling pathways, including inflammatory responses and regulatory pathways [108]. Interestingly, thromboxane A2 has been implicated in promoting neurite outgrowth in rat cortical neurons [77, 109]. In our system, levels of 2,3-dinor-TXB2 were reduced following DOX exposure in co-culture conditions. Inhibition of metabolic pathways associated with 2,3-dinor-TXB2 reduced neurite outgrowth, indicating that this signaling axis is important to the process of neurite outgrowth in our model.

### Human iPSC Model Supports Cardiac-induced Neuronal Deficits

To extend our findings to a human model system, we generated a hiPSC conditioned media-based cardio-neuronal co-culture model. Applying conditioned media from hiPSC cardiomyocytes exposed to DOX caused a reduction in βIII-tubulin fluorescent intensity in hiPSC sympathetic neurons, supporting the translational relevance of our findings. βIII-tubulin is a neuronal cytoskeletal protein, and its immunofluorescence is commonly used to assess neurogenesis, neuronal integrity, and neurite outgrowth [73, 74, 110, 111]. Additionally, NE levels significantly decreased in hiPSC sympathetic neurons treated with conditioned media from DOX-exposed hiPSC cardiomyocytes, suggesting impaired sympathetic neuronal activity. Importantly, these effects were observed in the absence of direct cellular contact, further supporting a role for paracrine signaling in cardio-neuronal communication. Collectively, these findings suggest that cardiomyocyte injury alters paracrine signaling in a manner that impairs neuronal structural integrity and function in a human cell-based system.

### Limitations and Future Directions

The limitations of the study include the following. The use of immortalized cell lines, while well-established, may not fully recapitulate the behavior of primary adult cardiomyocytes (despite the excitation-contraction coupling) and neurons. We began to extend key observations to female hiPSC-based models to improve physiological relevance and provide a more human-relevant platform; additional work is needed to further replicate and expand these findings in males and in human tissues. While metabolomic profiling revealed changes in metabolite abundance, the specific cellular source of the metabolites within co-cultures was not directly determined. While pharmacological inhibitors used on co-cultures may have off-target effects, they provide valuable insights at the pathway level, and the observed changes in neurite outgrowth support a functional role for these metabolic pathways.

Future studies should focus on the validation of these findings in vivo, in new approach methodologies, and on probing the mechanisms connecting cardiac metabolism to neuronal growth. Overall, this work identifies novel metabolic factors as critical mediators of cardio-neuronal communication, providing a foundation for future studies exploring therapeutic strategies for cardiac-induced neurotoxicity.

## Supporting information

Supplemental File Sloan et al.

## Acknowledgments

The human iPSC line was obtained from Joseph C. Wu, MD, PhD, at the Stanford Cardiovascular Institute, funded by NHLBI BhiPSC-CVD 75N9202D00019. We thank Dr. Wu for the support.

## Consent for publication

All authors approved the final manuscript before submission.

## Competing interests

The authors declare no competing interests.

## Funding

We thank the funding support to Dr. Viswanathan Rajagopalan from the New York Institute of Technology College of Osteopathic Medicine at Arkansas State University and the Arkansas Biosciences Institute (the major research component of the Arkansas Tobacco Settlement Proceeds Act of 2000). The funders had no role in study design, data collection and analysis, decision to publish, or preparation of the manuscript.

## Author Contributions

O.S.: Conceptualization, Methodology, Validation, Formal Analysis, Investigation, Writing—Original draft preparation, Writing—Reviewing and Editing, Visualization. S.O.: Methodology, Validation, Formal Analysis, Investigation. V.R.: Conceptualization, Methodology, Resources, Writing—Reviewing and Editing, Visualization, Supervision, Project administration, Funding acquisition.

## Notes

### Competing Interest Statement

The authors have declared no competing interest.

