## Supplemental File Sloan et al. for "Novel Cardiometabolic Factors Regulate Neurite Outgrowth in Cancer Chemotherapy-Induced Cardiotoxicity"

**Suppl. Figure 1: Dose-dependent Cardiotoxicity of Anti-cancer Agents**

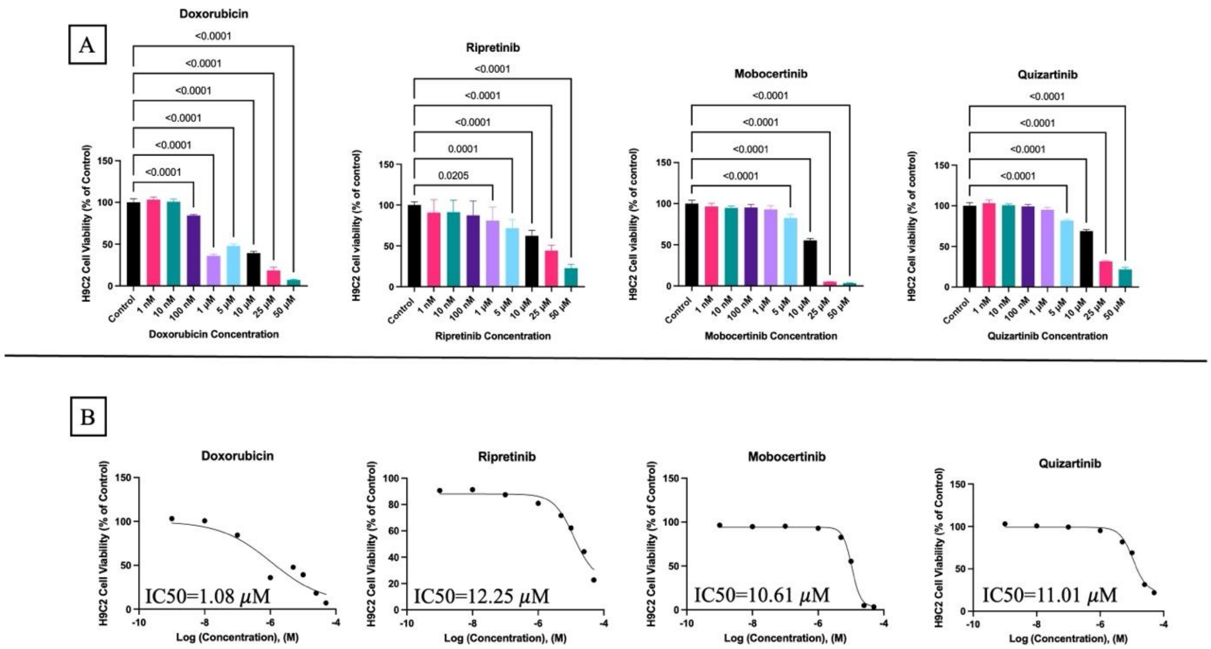


**Figure S1: (A)** Reduction in H9C2 cell survival following 24-hour treatment with (left to right) doxorubicin, ripretinib, mobocertinib, and quizartinib. Data is represented as means $\pm$ standard deviation (n=4). Data was analyzed via One-Way ANOVA and Tukey's post hoc test with GraphPad Prism. **(B)** Representative IC50 curves for treatment with (left to right) doxorubicin, ripretinib, mobocertinib, and quizartinib on H9C2 cells. IC50 values were determined by log-transforming molar concentrations and fitting the data using a four-parameter variable slope dose-response model in GraphPad Prism.

**Suppl. Figure 2: TOH Expression in DOX-exposed Co-cultures Compared to Controls**

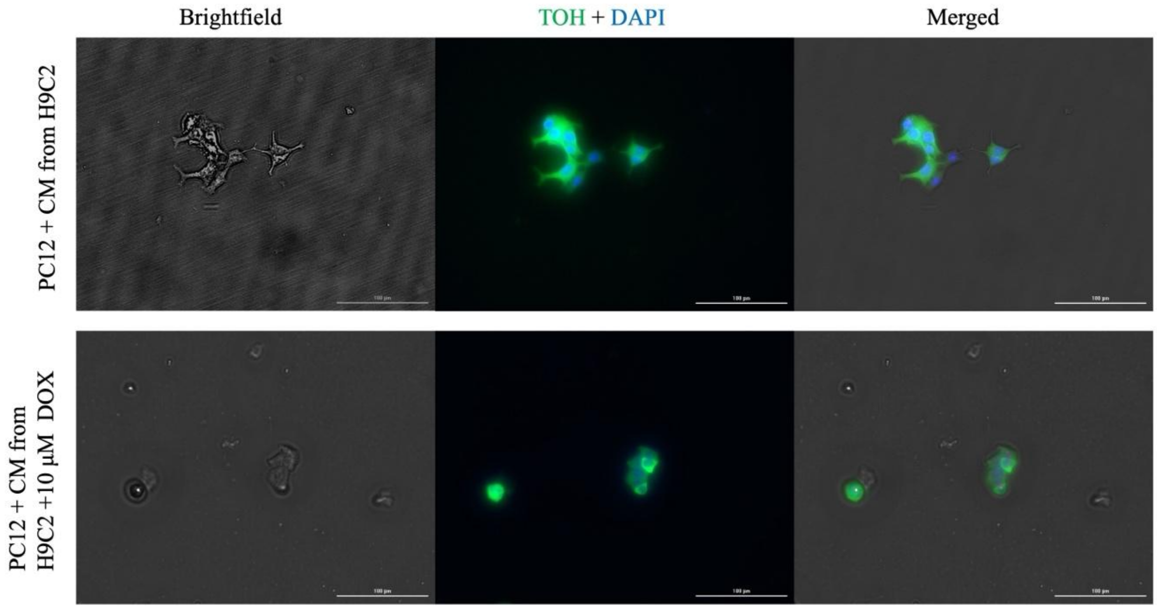


**Figure S2:** Immunofluorescent and brightfield images of control indirect co-cultures (PC12 + conditioned media [CM] from H9C2) (top panel) and 10 μM DOX-exposed indirect co-cultures (bottom panel) stained with tyrosine hydroxylase (TOH, neuronal marker, green) and DAPI (nuclear stain, blue). Scale bar = 100 microns; images were captured at 20x magnification.

**Suppl. Figure 3: Characterization of hiPSC-derived Cardiomyocytes and Sympathetic Neurons**

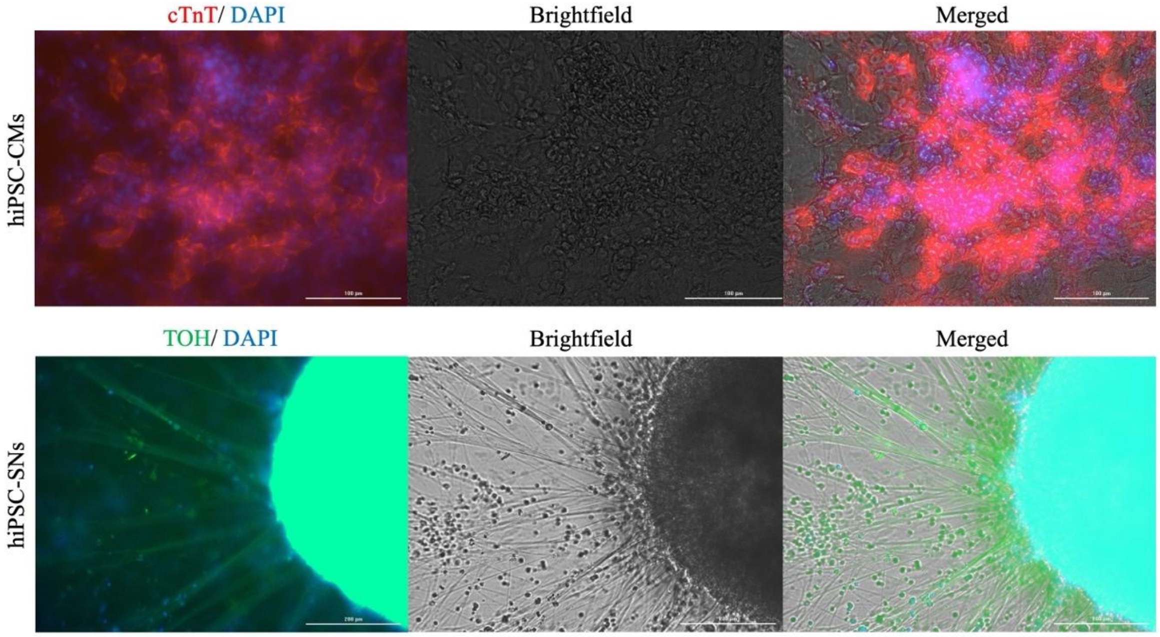


**Figure S3:** Representative immunofluorescent and brightfield images of hiPSC-derived cardiomyocytes (hiPSC-CMs) stained for cardiac troponin T (cTnT, red) (top panel) and hiPSC-derived sympathetic neurons (hiPSC-SNs) stained for tyrosine hydroxylase (TOH, green) (bottom panel). Cells were also stained for DAPI (nuclear stain, blue). Scale bars = 100 microns for hiPSC-CMs and 200 microns for hiPSC-SNs. Images were captured at 20x and 10x magnification, respectively.
